# Reanalysis of Bels et al. (2025) given complex anthropogenic hybridization in *Iguana delicatissima* and intraspecific diversity in *Iguana iguana*

**DOI:** 10.64898/2026.03.19.713079

**Authors:** Matthijs P. van den Burg, Julian Thibaudier

## Abstract

Understanding behavioral differences between hybridizing non-native and closely related endangered species could aid conservation management. In Zoology volume 169, Bels et al. (2025) presented first data on display-action-patterns (DAP) of native *Iguana delicatissima*, and compared those with non-native iguanas from islands of the Guadeloupe Archipelago, Lesser Antilles. Here, we address concerns about their study design and analytical methodology considering published literature on hybridization and reproductive seasons, and especially missing behavioral metadata preventing identifying and adjusting for potential confounding factors. Despite these limitations, we reanalyze their data given our proposed corrections, primarily a literature-informed adjustment of their two-level ‘species’ category through a site-level inference of hybrid status. Additionally, we utilize online videos to compare DAPs of non-native iguanas in the Archipelago with those from their populations of origin in South American. Significant differences in DAP characteristics among ‘species’ categories (native *I. delicatissima*, non-native iguanas, and hybrids) show that the variation in DAPs is more complex than suggested. Thus, highlighting the presence of behavioral variation among subpopulations within widely hybridizing iguanid populations, which has been poorly studied. Additionally, our preliminary DAP comparison between non-native iguanas and two mainland South American mitochondrial clades of *Iguana iguana* suggests that Guadeloupe non-native iguanas retained their original DAP. We show that the original dataset contains site-, individual-, and potentially context-dependent variation that cannot be assessed nor correct for, thus hampering considerations of the original hypotheses. Our effort will contribute to study design and methodology of future studies on iguanid behavior, especially in anthropogenic hybridization contexts.

## 1. Background

Invasive alien species (IAS) are a major global driver of biodiversity loss through a range of processes like predation, competition, parasite transmission, and disruption of reproductive interactions (e.g., Butchart et al., 2010; Vilà et al., 2011; Lymbery et al., 2014). In cases where IAS encounter closely related native species, behavioral interactions may be particularly disruptive. Similarities in communication and mate-recognition systems can facilitate heterospecific courtship, reproductive interference, and hybridization (e.g., Ord et al., 2001; Chapple & Wong, 2016; Galoyan et al., 2019). Understanding how behavior shapes interactions between IAS and endangered native species, is therefore important for predicting the outcomes of novel species interactions and their consequences for biodiversity conservation.

Native *Iguana* populations in the Caribbean Lesser Antilles comprise either *Iguana delicatissima* (Anguilla to Martinique, except on Saba and Montserrat; van den Burg et al., 2018a), or members of the *Iguana iguana* species complex (Saba, Montserrat, St. Lucia, St. Vincent and the Grenadines, and Grenada; Breuil et al., 2019; van den Burg et al., 2026). Additionally to these native populations, non-native iguanas (all part of the *I. iguana* complex) from different geographic origins have become established on most islands in the region. Genetic data shows these come from at least French Guyana, Curaçao, Colombia, and multiple countries in Central America (Stephen et al., 2013; Vuillaume et al., 2015; van den Burg et al., 2018b, 2023; Breuil et al., 2019; Pounder et al., 2020). Their presence has resulted in hybridization with native iguanas (e.g., Vuillaume et al., 2015), which has become the current most pressing threat to the survival of remaining native populations, whilst several have already become extinct; populations of *I. delicatissima* on St. Martin/St. Maarten, Barbuda, Antigua, and St. Kitts and Nevis (van den Burg et al., 2018a).

Since around 1860, multiple extra- and intra-regional incursions of non-native iguanas on Lesser Antillean islands have led to a widely varying status of currently present iguana populations (for pre-2008 incursions see Vuillaume et al., 2015; for post-2008 incursions see Breuil et al., 2019; van den Burg et al., 2020, 2023; Debrot et al., 2022; Angin & Guiougou, 2023). Concerning islands native for *I. delicatissima*, current populations either are still free of hybridization (e.g., Îles de la Petite Terre), have recent and ongoing hybridization (e.g., Dominica), have wide-spread hybridization with only few remaining pure *I. delicatissima* (e.g., Basse-Terre and Grande-Terre; Angin, 2017), or completely consist of non-native or introgressed individuals (e.g., Les Saintes and St. Martin).

Despite the region-wide decline of native *Iguana* populations in the Lesser Antilles, only few studies have assessed population dynamics and life-history characteristics of iguana populations where hybridization between native, non-native and hybrid iguanas takes place. Van den Burg et al. (2025) recently showed that hybrid iguanas seem to strongly contribute to genetic swamping through larger clutch sizes compared to native iguanas. Behavioral differences between native and non-native iguanas were not assessed prior to Bels et al. (2025), who analyzed patterns of head bob display. Beyond the first behavioral study in a hybridization setting, Bels et al. (2025) also provides the first assessment of head bob display action patterns (DAP) for *I. delicatissima*.

Head bob display in *Iguana iguana* is a research topic of sporadic nature (see Yan, 2014), since prior to Bels et al. (2025), reports on this topic originate mostly from the 20^th^ century and are limited to descriptive analyses of data from a few populations (Müller, 1972; Distel, 1978; Hazlett, 1980; Distel and Veazey, 1982; Dugan, 1982a,b; Rodda, 1992; Phillips, 1995). However, currently, we are aware of a high degree of geographically structured variation within this species complex (Stephen et al., 2013; van den Burg et al., 2026). Beyond potential intra- and interspecific differences in head bob display (see data from Dugan, 1982 and Distel and Veasey, 1982 in Table 1 of Bels et al., 2025), display behavior in reptiles is known to be dependent on other factors, including social and reproductive context but also temperature (see for examples Ferguson, 1977; Dugan, 1982a; Rodda, 1992; Rossi et al., 2023). Exploring differences in intra- or interspecific variation in display behavior should thus recognize these potential confounding variables.

**Table 1.**
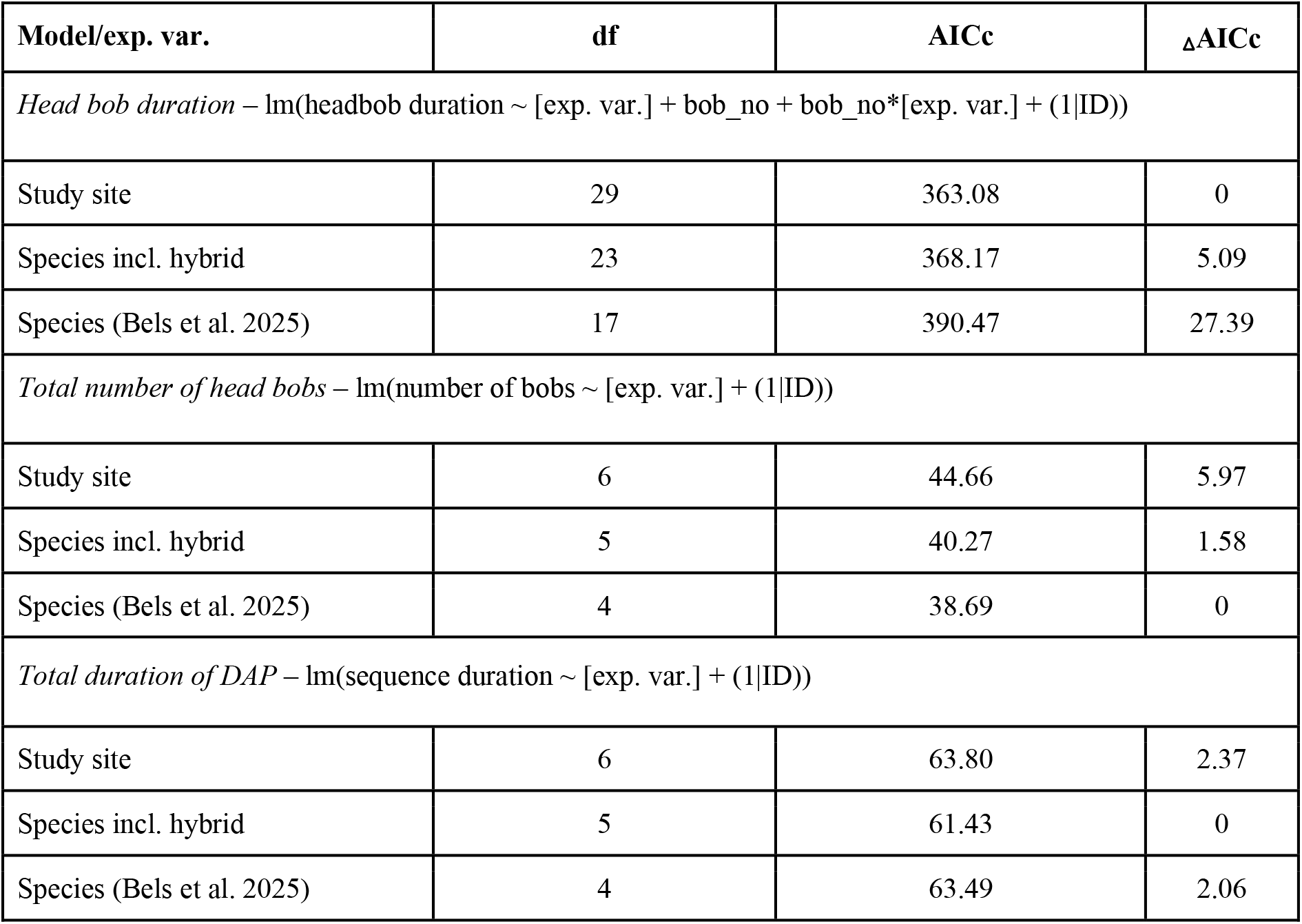
Comparison of model statistics (Akaikes Information Criterion modified for small sample sizes, AICc, and the difference between AICc scores, ΔAICc) between those presented by Bels et al. (2025) and in this manuscript. Abbreviations are explanatory variable (exp. var.), individual (ID), and head bob number (bob_no).

Given the renewed attention towards comparative behavior studies by Bels et al. (2025), we want to address the complexity of such comparisons in light of hybridization, temporal changes in behavior, and intraspecific variation. Our manuscript aims to 1) reclassify site-level admixture status and reassess DAP differences; and 2) compare non-native Guadeloupe DAPs to source-population (mainland) DAPs via video surveys.

## 2. Methods

Bels et al. (2025) described their study design as a comparison of DAPs isolated to those performed in a “challenge context”, a context in which congeners are present (Carpenter & Ferguson 1997). The Bels et al. (2025) dataset does not include information on additional social context of each recorded DAP, such as how many female and male iguanas were present or their distances from the displaying male. Bels et al. (2025) state that they removed “DAPs that were followed by an attempt to copulate to confine observations to the assertion context”. However, based on the information available in the dataset, it is difficult to determine whether this procedure was sufficient to distinguish DAPs performed in a challenge context from those occurring in other social contexts.

To build their dataset, Bels et al. (2025) recorded videos of behavioral display from *I. delicatissima* during February until the end of April (unknown year, although the Result section mentions February–March of 2013 and 2014). Since *I. delicatissima* on La Désirade lay eggs between June–mid August (van den Burg et al., 2018a) and because iguanas lay eggs about six to eight weeks after copulation (van Marken Lichtenbelt & Albers, 1993; Frye, 1995), the period of February–April partially coincides with the copulation period. Additionally, the videos of iguanas in the non-native iguana category were recorded during February of 2013 and between February–April of 2014, and Bels et al. (2025) indicated that according to Breuil (2002) that period falls before the reproduction period. However, we currently know that the reproduction periods of non-native iguanas and *I. delicatissima* on Guadeloupe and La Désirade are not identical (see Figure 17 in Breuil et al., 2019), with February–April partially falling within the nesting period of non-native iguanas, and partially falling in the copulation period of *I. delicatissima*. Therefore, it cannot be ruled out that aspects of display behavior changed during the study period due to progression of the reproduction season (Rodda, 1992), biasing the dataset. In addition to these potential effects of reproductive timing, the general social setting might have affected the dataset too, since the authors describe how *I. delicatissima* displayed within stereotyped displays at lekking sites, whilst non-native iguanas displayed more randomly. We note that metadata such as date, time of day, temperature, group size, and social context are not available for individual video recordings in the Bels et al. (2025) dataset. Consequently, it is difficult to determine whether and to what extent these potential confounding variables may have influenced their findings.

Bels et al. (2025) summarize that about half of the genetically assessed iguanas from the Guadeloupe Archipelago concerned hybrid or introgressed individuals (Vuillaume et al., 2015). They additionally note that whilst some native *I. delicatissima* remain present on Basse-Terre and Grande-Terre (Angin, 2017), most iguanas are not pure *I. delicatissima*. After addressing these points, Bels et al. (2025) state that they “used the term of non-native *Iguana* for specimens studied in the main islands of the Archipelago”. Here, we reason that treating all iguanas on Basse-Terre and Grande-Terre as one entity is not in line with the cited references. Namely, considering known locations of ongoing hybridization between pure *I. delicatissima* and non-native iguanas (defined as pure non-native, hybrid or introgressed; see figure 3 in Angin, 2017; B. Angin pers. comm. 2026), the field sites where Bels et al. (2025) studied DAPs seemingly represent sites with known or suspected hybridization/introgression. Specifically, 1) the site at Gosier (Grande-Terre) lies within a region from where only non-native iguanas (B. Angin pers. comm. 2026) are known, 2) the site at Malendure (Basse-Terre) lies near an area with known ongoing hybridization, and 3) the site at Carangaise (Basse-Terre) lies within an area with known hybridization (Angin, 2017; Bels et al., 2025). Given these references, we argue that an assessment that regards hybrid iguanas as a separate group is better justified compared to the general assignment as substantiated by the references in Bels et al. (2025). Hence, we argue that the ongoing presence of pure *I. delicatissima* and hybridization of these with non-native iguanas near or at the Malendure and Carangaise sites should be acknowledged within the “species” assignment. To address this, we added “hybrid” as a third category, to the two original groups, “*Iguana delicatissima*” and “non-native *Iguana*”, of Bels et al. (2025). We assigned the Colibri site (CO) as “*Iguana delicatissima*”, the Malendure (MA) and Carangaise (CA) sites as “hybrid”, and the Gosier site (GO) as “non-native”. As the Bels et al. (2025) dataset does not provide any information on the genetic make-up of each individual (such as which generation of hybrid), nor genetic information, this adjusted species variable should be regarded as a site-level inference instead of a genetic-level inference.

Here, we reanalyzed the data of Bels et al. (2025) (as provided in their supplementary material files) including the new “hybrid” category. Within the *R* environment (R Core Team, 2024), we utilized the same methodological approaches for data transformation and statistical tests as Bels et al. (2025) to assess for differences among three DAP characteristics: the duration of individual head bobs, the number of head bobs per DAP, and the total duration of the DAP. In short, we built linear models using log-transformed response variables and the three-category species variable as the explanatory variable through the *Lme4* package (Bates et al., 2015). However, considering the likely different stages of hybridization and introgression among the MA, CA, and GO sites, we also built linear models where we replaced “species” with “site” as an explanatory variable. Since the species and site variables are confounded, it was not possible to assess these within the same model. Subsequently, we compared Akaikes Information Criterion modified for small sample sizes (AICc) scores from the *MuMIn* package (Bartoń, 2026) among the two models (“species” or “site”), to select the model with the best fit per assessed DAP characteristic. For the best fitted model, we performed type III F-tests and Likelihood ratio tests to test fixed and random effects respectively, similar to Bels et al. (2025). We simulated residuals with the *DHARMa* package (Hartig, 2025). Subsequently, we performed pairwise comparisons using the *emmeans* function (Russell, 2024) with the default Tukey setting to test for significant differences among categories of the explanatory values (“species” or “site”) for all three response variables. Effect sizes were calculated using ‘Cohens *d*’ from the same package. We did not apply a family correction, since our aim is to follow the methodology of Bels et al. (2025) in which DAP characteristics are tested separately, and therefore this would result in unnecessary loss of statistical power (Carpenter and Ferguson, 1977; Carpenter, 1978; Stegmann, 2013; Barnett et al., 2022). Additionally, we compared the AICc scores of our models with AICc scores of models with only the original, binary species category of those of Bels et al. (2025). The robustness of the selected model was visually evaluated by comparing its output with that of the other two models. We provide our *R* code, see File S1in the Supporting Information section of this paper, and the data are available in Data File S1 and S2. We encountered some discrepancies between the supplementary files, data and figures in Bels et al. (2025), which we summarized in our File S2 to aid others in understanding the supplementary files we used.

Considering data visualization, we could not create 100% identical boxplots (e.g., equal number of outliers) as these details are not included in *R* code of Bels et al. (2025) (see supplementary materials of Bels et al., 2025). We compared their boxplots with those created with default settings using the *boxplot* and *geom_boxplot* functions (Wickham, 2016; R Core Team, 2024). We generated the boxplots presented here using the default *geom_boxplot* function.

Our reanalysis clarifies that the position of successive head bobs per DAP as visualized in Figure 2 of Bels et al. (2025) were not equal between both species groups. Namely, whilst head bob numbering of *I. delicatissima* are presented in their respective columns (left-ordered), these same data for non-native *Iguana* were right-ordered, meaning that only data for DAP sequences with eight head bobs are represented in the first column of the non-native *Iguana* subfigure (Figure 2 of Bels et al., 2025); the eighth column thus represents the last head bob for each DAP sequence. Here, we adjusted these data so that durations of identically numbered successive head bobs can be compared among the different groups, see our Figure 2. We highlight that this is an important characteristic regarding the variation in display behavior within a population and include an overview figure of these data (Figure 1).

**Fig. 1.**
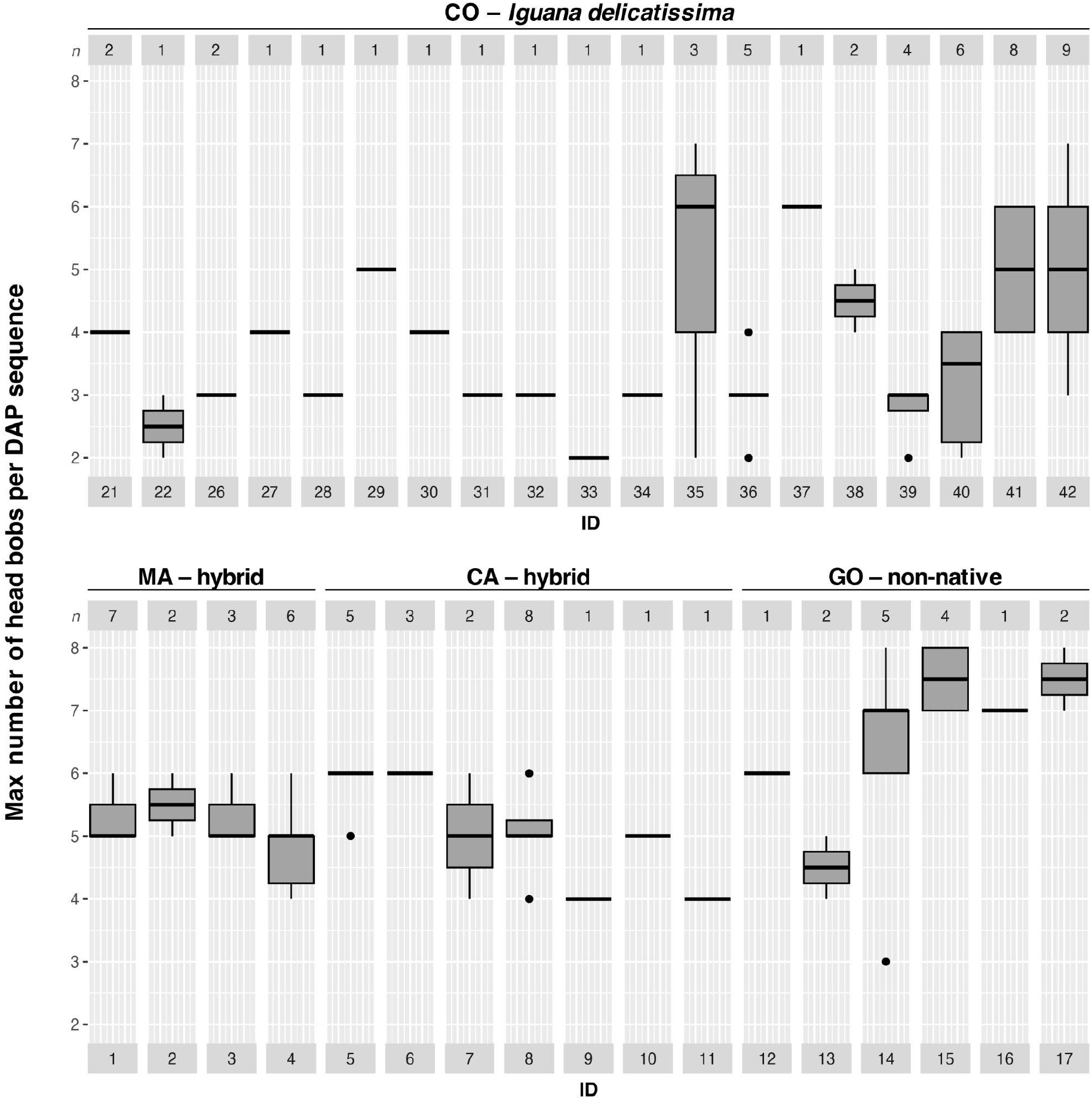
Variation in number of max head bobs per DAP per individual (ID) across the different sites. Grey boxes represent the ID code for each iguana (lower) and the number of DAPs the corresponding boxplot represents (upper).

**Fig. 2.**
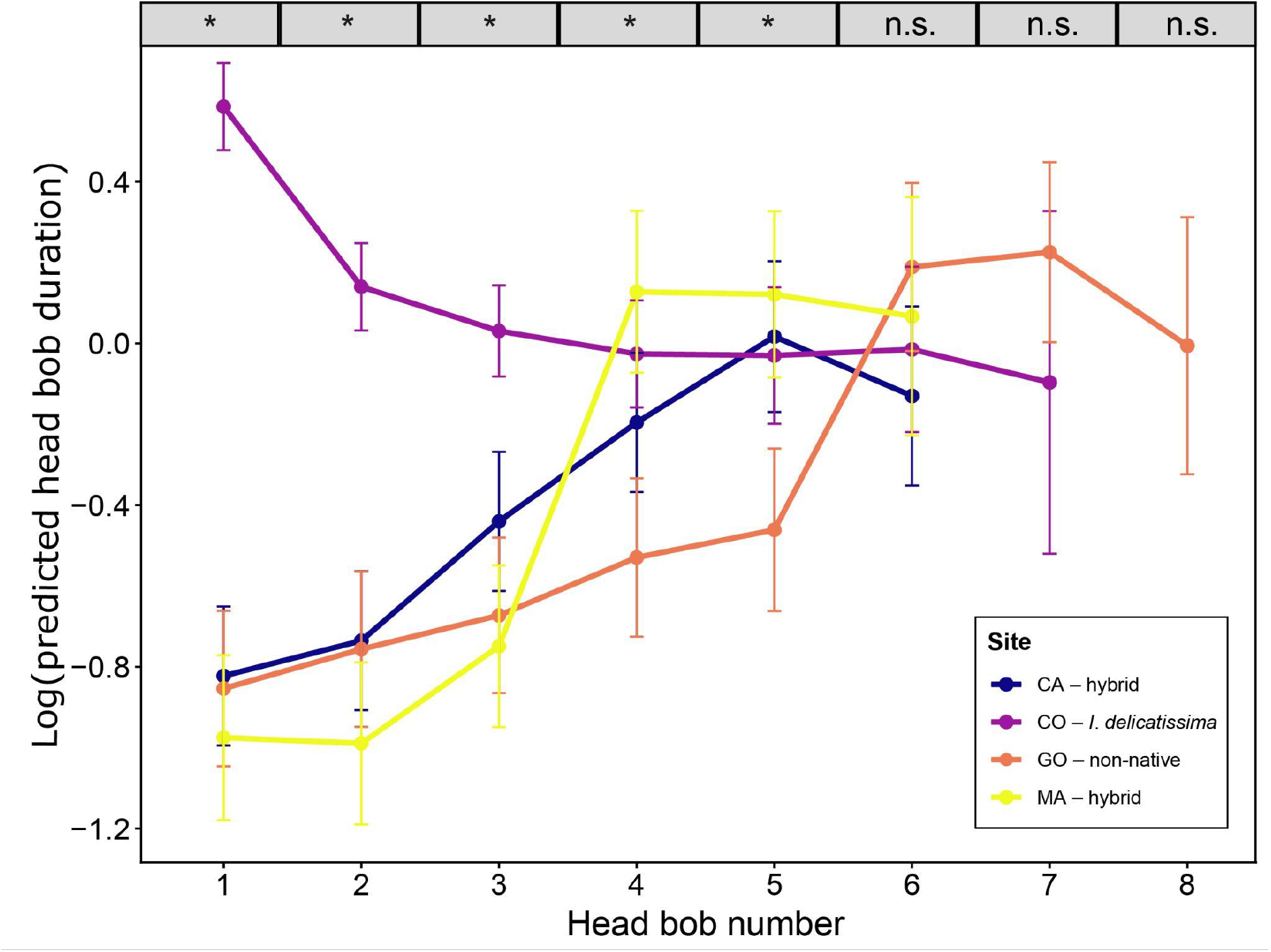
Log transformed prediction of head bob duration, for iguana populations at Colibri (CO; purple), Carangaise (CA; blue), Malendure (MA; yellow), and Gosier (GO; orange). Head bob numbers that showed significant differences among at least two of the sites are indicated with an asterisk, head bob numbers without significant differences with n.s.

**Fig. 3.**
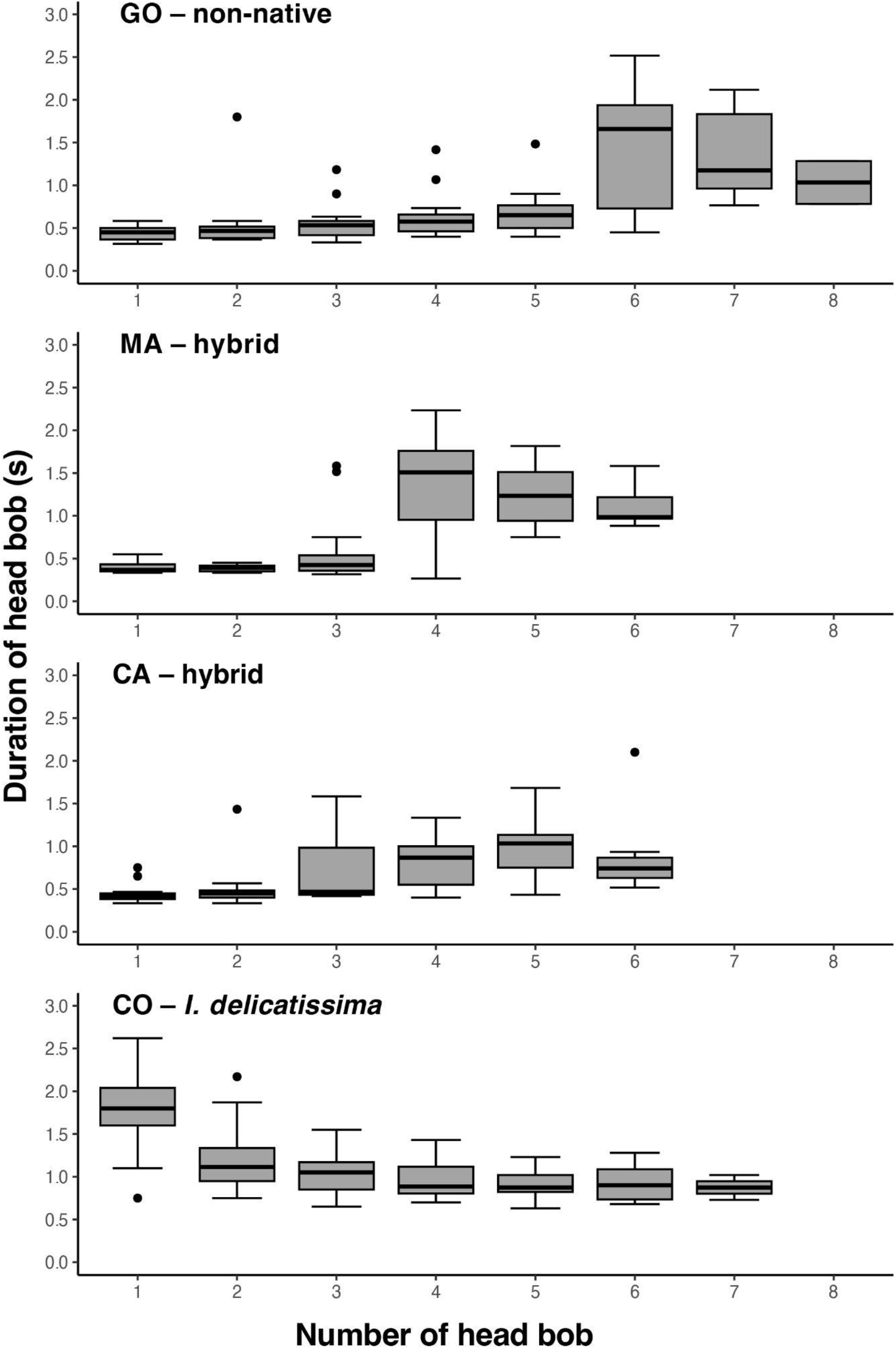
Variation in the duration of head bobs across the study sites. Each head bob sequence starts at head bob number 1, in contrast to Bels et al. (2025).

Lastly, Bels et al. (2025) noted the importance of comparing DAP characteristics of the non-native iguanas on Guadeloupe with those from mainland French Guyana *I. iguana* populations, which are the source of non-native iguanas on Guadeloupe (Vuillaume et al., 2015). The authors state that those data were not available and instead utilized data from two studies on native *I. iguana* from the South American mainland; Panama (Dugan, 1982a) and Colombia (Distel and Veasey, 1982). Whilst non-native iguanas on Guadeloupe begin their DAP with at least a single short head bob and end with longer head bobs, the DAP from iguanas in Panama (Dugan, 1982a) and Colombia (Distel and Veasey, 1982) start with longer head bobs and finish with shorter head bobs. In their discussion, Bels et al. (2025) discuss several hypotheses regarding the potential retainment or evolution of DAP of non-native iguanas once these hybridized and introgressed with *I. delicatissima*. Here, we take this geographic variation of *I. iguana* into account, by performing a preliminary comparison of DAPs of non-native iguanas with those of the similar mtDNA clade, which we show is a valuable addition to the analyses of Bels et al. (2025).

Bels et al. (2025) compare the position of short and long head bobs between the non-native iguanas on Guadeloupe with native mainland populations. Therefore, to compare this basic DAP characteristic between non-native iguanas from Guadeloupe and their original population, we searched for freely accessible online videos showing green iguanas from mainland locations of countries where Clade II and/or IV (sensu van den Burg et al., 2026) are present (Clade IIA and IIB sensu Stephen et al., 2013; van den Burg et al., 2021): Costa Rica, Panama, Colombia, Venezuela, Guyana, Suriname, French Guyana, Brazil, Peru and Ecuador. We acknowledge the difficulty of collating life history data from wide-range species and therefore used keywords in English, Spanish, French and Portuguese (van den Burg and Kaiser, 2026). We used the following key string: (iguana verde OR iguana OR Iguana iguana OR iguane vert OR green iguana) AND (Costa Rica OR Panama OR Colombia OR Venezuela OR Guyana OR guiana OR Suriname OR French Guyana OR Guyane française OR Brazil OR Brasil OR Peru OR Ecuador OR Acacias OR amazonas OR amazon OR amazona OR Manaus OR Pantanal OR mato grosso OR Tocantins OR Piaui OR Natal OR nordeste OR nordestina). We performed searches between 20–27 January 2026 on the following platforms: Google, Tiktok, Youtube, and Instagram. Videos that included head bobbing male *I. iguana* were retained, for which we assessed whether longer head bobs (similarly to “unit 1” in Dugan, 1982a) occurred anteriorly or posteriorly to shorter head bobs; we did not create DAP diagrams. We used location data and visual assessment of each iguana to make sure retained videos were all from different iguanas. We caution that this is a preliminary assessment since we were not able to place the videos in the context of behavior nor that of the reproductive season.

## 3. Results

### 3.1. Display behavior

For 36 individuals data on the number of head bobs per DAP was available, of which 21 individuals had data from more than a single DAP (Figure 1). Of these 21, the number of head bobs per DAP was identical across their DAPs for 14% (three individuals). The highest number of DAPs of an individual was nine, and six individuals had data from more than five DAPs.

### 3.2. Head bob duration

Based on AICc comparison, the model including site (AICc = 363.08) was better supported compared to the model including species (AICc = 368.17). The duration of individual head bobs significantly differed between sites (F_4,35_ = 40; P < 0.001), with the head bobs of the *I. delicatissima* population at CO being longer compared to other iguanas at the other sites (see Table S1 for effect sizes). Furthermore, there was a significant interaction effect between site and head bob number (F_16,453_ = 29; P < 0.001) with head bob duration of *I. delicatissima* at CO decreasing during the sequence, and head bob duration of iguanas from the other three sites increasing during the sequence. More specifically, the pairwise comparison test revealed the first three head bobs to be longer for *I. delicatissima* compared to hybrid and non-native groups (P < 0.001), but with no difference between hybrid and non-native groups (Figure 2). The fourth head bob is longer in *I. delicatissima* and hybrids at MA compared to non-native groups at GO (P < 0.001), but not longer for hybrids at CA compared to non-native groups at GO. The fifth head bob in the DAP of non-native groups is shorter compared to *I. delicatissima* and hybrids (P < 0.008; for full table of pairwise comparisons and effect sizes, see Table S1). From head bob 6 onwards, results indicated there were no significant differences between species. Corroborating Bels et al. (2025), our analysis also substantiated a significant random effect due to different head bob duration among individual iguanas (X^2^ = 41, df = 1, P < 0.001). Figure 3 provides an overview of the variation in head bob duration per site, with all head bob sequences starting at the first column of the x-axis.

Significant differences for head bob 1–3: *I. delicatissima* longer than all others; head bob 4: *I. delicatissima* and hybrids at MA longer than the non-natives; and head bob 5: non-native shorter than all other iguanas.

### 3.3. Total number of head bobs

AICc indicated that including species (AICc = 40.27) in the model was better supported compared to site (AICc =44.66). Species had a significant effect on the number of head bobs per DAP (F_3,32_ = 510; P < 0.001), with a pairwise comparison test showing that DAP of *I. delicatissima* containing significantly (df = 32; P < 0.001 and df = 27; P <0.001, respectively) fewer head bobs compared to both the non-native (*d* = -2.413; CI 95% = -3.434 - -1.392and hybrid groups (*d* = -1.534; 95% CI = -2.327 - -0.741). No significant (df = 29; P = 0.2) difference between the number of head bobs of the hybrid and non-native groups was found (*d* = 0.879; 95% CI = -0.156 - 1.914) Corroborating Bels et al. (2025), our analysis also substantiated a significant random effect due to a different number of head bobs among individual iguanas (X^2^ = 16, df = 1, P < 0.001). Figure 4 provides an overview of the variation in head bobs per DAP per study site.

**Fig. 4.**
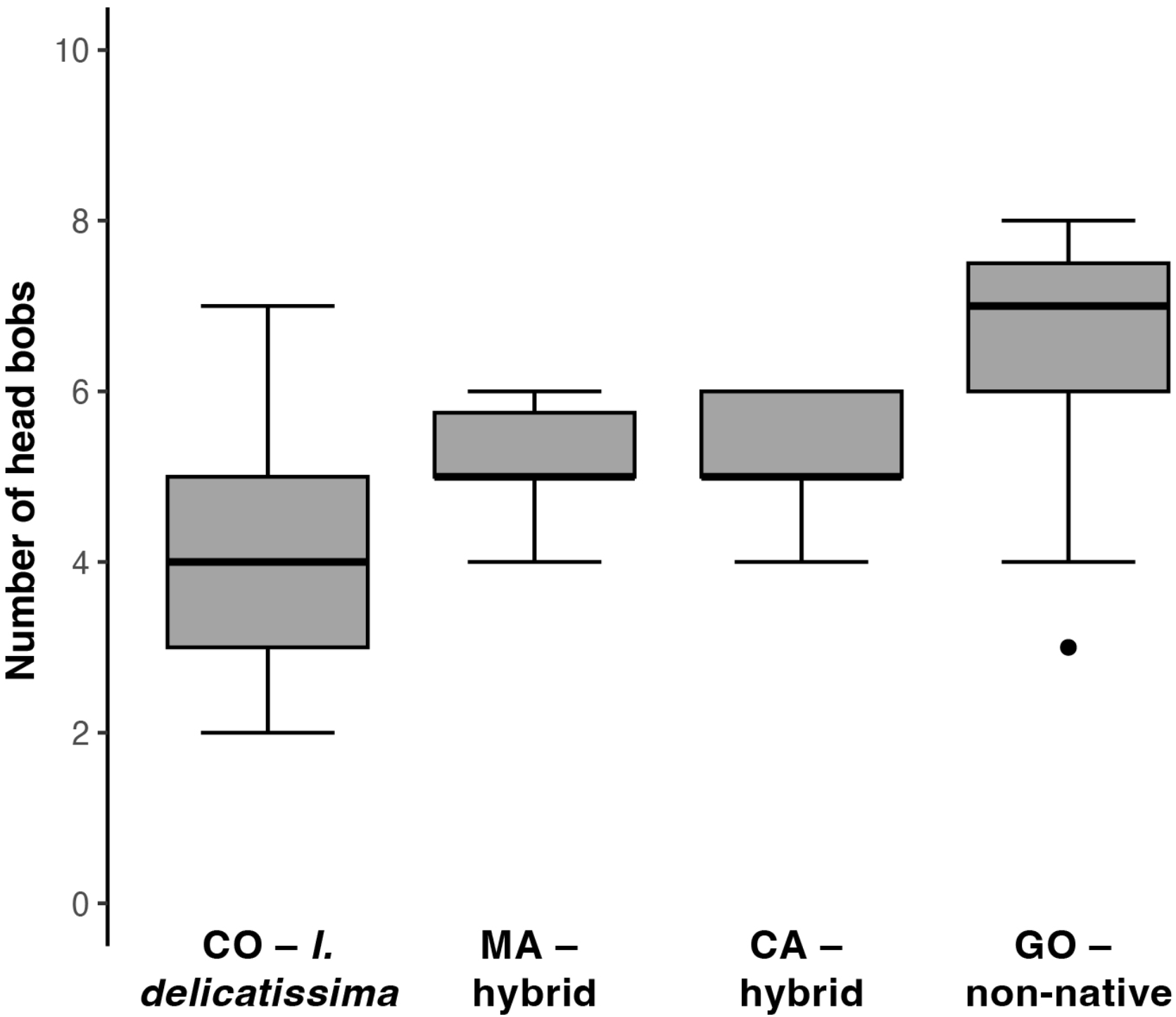
Variation in the number of head bobs per DAP across the study sites.

### 3.4. Total duration of head bobs

The AICc comparison indicated that the model including species (AICc = 61.43) was more supportive compared to the model including site (AICc = 63.80), albeit the ΔAICc is small (ΔAICc = 2.37). Species is a significant variable regarding the duration of head bob sequences (F_4,33_ = 575; P < 0.001), with the pairwise comparison showing that DAP duration differed between all species groups: *I. delicatissima* have a significantly (df = 32; P = 0.034 and df = 28; P < 0.001, respectively) longer DAP duration compared to the non-native (*d* = 1.43; 95% CI = 0.293–2.57) and hybrid groups (*d* = 2.91; 95% CI = 1.893–3.93), and hybrids have a significantly (df = 33; P = 0.036) shorter DAP compared to non-native group (*d =* 1.48; 95% CI = 0.296–2.66). Corroborating Bels et al. (2025), our analysis also substantiated a significant random effect due to a different duration of DAP among individual iguanas (X^2^ = 26, df = 1, P < 0.001). Figure 5 provides an overview of total DAP duration per study site.

**Fig. 5.**
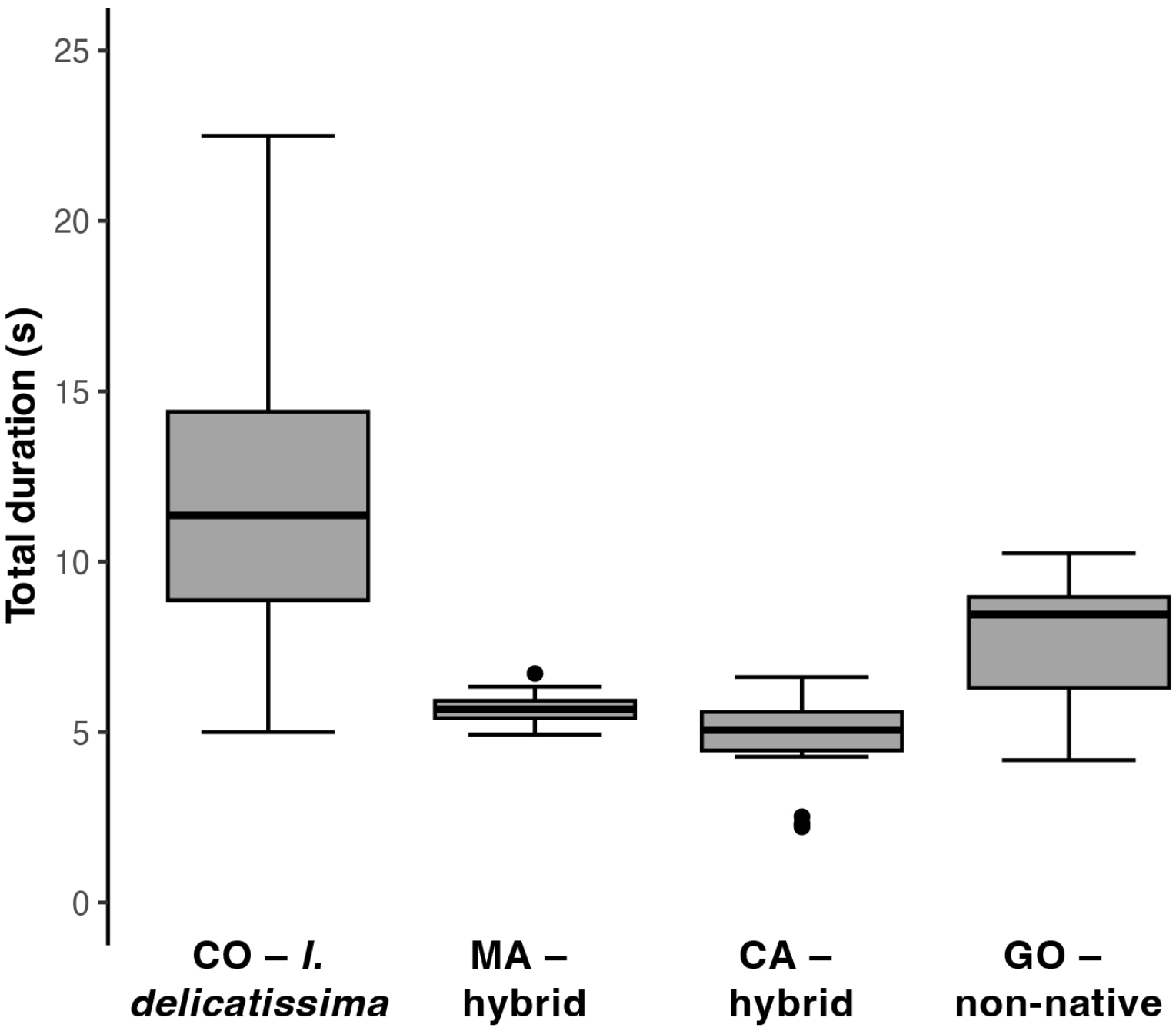
Variation in total duration of DAP across the study sites.

### 3.5. Model comparison with Bels et al. (2025)

Comparison of AICc scores among models generated here and by Bels et al. (2025) showed that the variable with the best model fit was different for each tested DAP characteristic. Namely, for the ‘head bob duration’ model the study site showed the best model fit, for the ‘total number of head bobs’ model the species variable using ‘native’ and ‘non-native’ following Bels et al. (2025) showed the best fit, and for ‘total duration of DAP’ the species variable including ‘hybrid’ showed the best model fit (Table 1).

### 3.6. Comparison between mainland *I. iguana* (South America) and non-native iguanas (Guadeloupe)

Our online survey identified a total of 23 usable videos: 11 from Clade II and 12 from Clade IV (see summary data and video links in Table S2). DAPs from localities within the range of Clade II (including Colombian areas west of the Cordillera Oriental) started with long head bobs that were followed with shorter head bobs, identical to data from Dugan (1982a) and Distel and Veasey (1982). In contrast, DAP sequences from localities in Clade IV (including Acacias, directly east of the Cordillera Oriental), southeast of the Andes, were consistently reversed starting with short head bobs before longer head bobs (see Table S2).

## 4. Discussion

Bels et al. (2025) provide an important first assessment of display behavior for *Iguana delicatissima* and compare that to non-native iguanas (unknown mix of pure non-native, introgressed, and hybrid iguanas) that form a major threat to this critically endangered species. Assessing behavioral aspects of this anthropogenic hybridization is important as it could provide clues about underlying reasons of the observed outcome, the eventual disappearance of the native *I. delicatissima* (van den Burg et al., 2018a). This first assessment and comparison, however, was built on a dataset and analytical methodologies that either cause biased results (e.g., equal sampling period but different reproduction season) or prevents assessing influences of confounding factors (e.g., absence of metadata like behavioral context). We identified limitations considering different aspects of the study design and analyses, namely: 1) small and uneven sample sizes; 2) the presence of individual-level variation, which was not reported; 3) confounding of site and category; 4) lack of metadata and control for date, time, and temperature; 5) lack of metadata for behavioral context whilst presence of observed differences were acknowledged; 6) absence of genetic information from studied individuals.

Here, we argue that the approach taken by Bels et al. (2025) did not fully capture the complexity of hybridization on Basse-Terre and Grande-Terre. Our reanalyzes of their data using a three-level species variable (including hybrids) shows that all three DAP characteristics are different between native compared to both the hybrid and non-native *Iguana* groups, corroborating Bels et al. (2025). In addition, we also showed that DAP characteristics differ between the hybrid and non-native iguana groups, namely the duration of the fourth and fifth head bob, as well as the total duration of DAPs. Thus, we argue that our methodology is a better representation of the natural situation in Guadeloupe, as also supported by a better model fit for two of the three models, albeit one of these by a relative low ΔAICc score (2.37), we consider this still informative given that AICc scores include a penalty for adding parameters. Additionally, we provide insights on the individual variation in the number of head bobs per DAP, which is an important descriptive display characteristic. Overall, our addition to the work by Bels et al. (2025) substantiates that studies on hybridization in iguanids should acknowledge the complexity of hybridization. Nonetheless, we acknowledge that the DAP of known F1 hybrids has not been assessed, and thus we are unaware how hybridization precisely affects changes in display behavior. We note that it would be important to study the behavior of individuals with genetically corroborated signatures, which ideally include F1 and later-stage introgressed individuals.

Does an iguana always head bob an equal number of times per DAP? We used the Bels et al. (2025) dataset to assess this important behavioral feature (see Figure 1), indicating a high degree of variation per individual. Considering only the 21 individuals with >1 recorded complete DAP, just 14% (two *Iguana delicatissima*, one animal from CA) had an equal number of head bobs across their recorded DAPs. This is corroborated by significant X^2^ values for all models, indicating significant variation in DAP characteristics among individuals. We note that Bels et al. (2025) only included DAPs that were complete, but no definition for a complete DAP was provided, nor an explanation of *in situ* conditions that led to individual iguanas not completing a DAP; for example, on Saba we have seen that males can pause or terminate DAPs when a new individual appears into their view (pers. obs., MP van den Burg). The variation in head bobs per DAP furthermore questions how representative the number of head bobs were of individuals from which only a single DAP was recorded. Does the observed intra-individual variation reflect different behavioral contexts? And how many DAPs per individual, and number of individuals, represent a population’s DAP phenotype? Given the considerable variation in DAP per individual observed here, future studies should assess this characteristic and consider including only those individuals from which a minimum set of DAPs were recorded, as well as providing more details on individual’s variation.

Our preliminary assessment of short and long head bob placement within a DAP using on-line accessible videos, suggests there is a difference between DAPs of male iguanas from Clades II and IV (sensu van den Burg et al., 2026). We note that although the four major mtDNA clades within the *I. iguana* species complex acknowledge its high intraspecific variation (Stephen et al., 2013), within-clade diversity is more continuous (van den Burg et al., 2026). Jointly, this justifies extrapolating data from Clade IV members (Suriname, Venezuela, Colombia, and Brazil) to male *I. iguana* from French Guyana, populations that are also part of Clade IV (van den Burg et al., 2026). A successive comparison of this DAP characteristic shows that the Clade IV individuals in our limited dataset and the non-native iguanas on Guadeloupe (Bels et al., 2025) both display long head bobs at the end of their DAP, suggesting that the latter might have retained their original display behavior. However, we note the descriptive nature of this comparison given the absence of metadata that are necessary to consider potential confounding variables (e.g., behavioral context, time, temperature, etc.). This finding should therefore be confirmed in a future assessment based on a robust dataset.

Here, we considered the methodological design and analyses performed by Bels et al. (2025), and conducted a reanalysis of their data. Our analyses identified variation across sites, individuals, and potentially social contexts within the dataset, highlighting the possibility that factors other than species identity may contribute to the observed patterns. Because these considerations also constrain our reanalysis, the available data do not allow us to draw definitive conclusions regarding the main research question of Bels et al. (2025). Likewise, they limit the extent to which the hypotheses discussed in their Discussion can be further evaluated using these data. We therefore complemented the findings of Bels et al. (2025) by highlighting methodological considerations that may be useful in informing future studies of iguanid behavior, especially in an anthropogenic hybridization context.

## CRediT authorship contribution statement

### Matthijs P. van den Burg

Conceptualization, Data curation, Investigation, Writing – original draft.

### Julian Thibaudier

Data curation, Formal analysis, Writing – original draft.

### Appendix A

Supplementary data associated with this article can found in the online version at …

### Data availability

All data are available in the Supplementary files, or Supplementary files of Bels et al. (2025).

