## Supplementary material for "Reanalysis of Bels et al. (2025) given complex anthropogenic hybridization in *Iguana delicatissima* and intraspecific diversity in *Iguana iguana*": File S1

\# Reconstruction of data analysis

#Bels, V. L., Brousse, C., Pelle, E., Guerlotté, J., Pierre, M. A., Kirchhoff, F., & Biro, P. A. (2025). Comparative display behaviour of the native Iguana delicatissima with the non-native Iguana in the Guadeloupe Archipelago (Lesser Antilles). Zoology, 169, 126239. DOI: https://doi.org/10.1016/j.zool.2025.126239

setwd('your working directory')

\# Packages

\# Models and analysis

library(lme4)

library(emmeans)

\# For plotting

library(ggplot2)

library(ggeffects)

library(viridis)

\# Data handling

library(dplyr)

\# Model evaluation

library(DHARMa)

library(lmerTest)

library(broom.mixed)

###########################################################

\#### Loading data, preparing data #########################

###########################################################

durbob <-read.csv("1dur_bob.csv", sep = ';')

#totbob <-read.csv("2tot_bobs.csv") # missing

totdur <-read.csv("3tot_dur.csv", sep = ';')

i_deli <- durbob$spp == "deli"

i_iguana <- durbob$spp == "iguana"

durbob <- durbob %>%

mutate(

species = case_when(

loc == "COLIBRI" ~ "Iguana delicatissima",

loc %in% c("MA", "CA") ~ "Iguana iguana x delicatissima",

loc == "GO" ~ "Iguana iguana",

TRUE ~ NA_character_

)

)

######### For figure 2, the head bob duration

\# Remove outliers

sum(is.na(durbob$dur_bob))

durbob$dur_bob[durbob$dur_bob > 10] <- NA

sum(is.na(durbob$dur_bob))

\# Iguana iguana head bob number order is turned around in the datasheet. The following lines assign a individual sequence number to all the sequences. This will later help turn around the Iguana iguana head bob number.

\# Add sequence number

durbob$seq_no <- NA # initialize

durbob$seq_no[i_deli] <- cumsum(c(1, diff(durbob$bob_no[i_deli]) < 0)) # deli: normal sequence (increasing when hb_no decreases)

durbob$seq_no[i_iguana] <- 52 + cumsum(c(1, diff(durbob$bob_no[i_iguana]) > 0)) # iguana: reverse sequence (increasing when hb_no increases)

\# Now the actual turning around of the head bob numbers

durbob$bob_nocor[i_iguana] <- ave(

durbob$bob_no[i_iguana],

durbob$seq_no[i_iguana],

FUN = function(x) rank(-x, ties.method = "first")

)

durbob$bob_nocor[i_deli]<- durbob$bob_no[i_deli]

\# Make bob number a factor, for fixed effect in model

durbob$bob_nocor <- as.numeric(durbob$bob_nocor)

durbob$bob_no2 <- as.factor(durbob$bob_nocor)

######### For figure 3, number of head bobs per sequence. This datasheet is missing from supplementary material, so reconstructed with 1dur_bob.csv

\# To group together, add sequence number to individual sequences. This is also loaded in previous lines

#durbob$seq_no <- NA # initialize

#durbob$seq_no[i_deli] <- cumsum(c(1, diff(durbob$bob_no[i_deli]) < 0)) # deli: normal sequence (increasing when hb_no drops)

#durbob$seq_no[i_iguana] <- 52 + cumsum(c(1, diff(durbob$bob_no[i_iguana]) > 0)) # iguana: reverse sequence (increasing when hb_no rises)

\# Group the data to recreate 2tot_bob.csv

totbob <- durbob %>%

group_by(spp, species, loc, ID, seq_no) %>%

summarise(

nr_bobs = n (),

seq_duration = sum(dur_bob),

.groups = "drop"

)

totbob %>% count(species)

########## For figure 4

\# Add hybrid status to the datasheet

totdur <- totdur %>%

mutate(

species = case_when(

loc == "COLIBRI" ~ "Iguana delicatissima",

loc %in% c("MA", "CA") ~ "Iguana iguana x delicatissima",

loc == "GO" ~ "Iguana iguana",

TRUE ~ NA_character_

)

)

totdur %>% count(species) #four hybrid datapoints missing, compared tot totbob

###########################################################

\#### Fig. 2: Head bob duration ############################

###########################################################

\# Data is skewed, log transform

hist(durbob$dur_bob)

durbob$dur_boblog = log(durbob$dur_bob)

hist(durbob$dur_boblog)

\# What Bels et al. 2025 did:

bfit2 <- lmer(dur_boblog ~ 0 + spp + bob_no2 + bob_no2:spp + (1|ID), data = durbob)

anova(bfit2) #result from Bels: spp: F2, 131 = 50.9, P < 0.0001 & spp*bob: F7,472 = 16.9, P < 0.0001

ranova(bfit2) #result from Bels: LRT, X2 = 77.6, df = 1, P < 0.0001

summary(bfit2)

\# What we do:

fit2 <-lmer(dur_boblog ~ 0 + loc + bob_no2 + bob_no2:loc + (1|ID), data= durbob)

fit21 <-lmer(dur_boblog ~ 0 + species + bob_no2 + bob_no2:species + (1|ID), data= durbob)

AIC(fit2, fit21, bfit2)

\# Results

anova(fit2)

ranova(fit2)

summary(fit2)

\# Post hoc model evaluation and predictions. Adapt to desired model

pr <- ggpredict(fit2, c("loc"))

plot(pr)

z <- augment(fit2)

head(z)

hist(z$.resid)

plot(DHARMa::simulateResiduals(fit2))

\# Pairwise comparison test (optional)

emm2 <- emmeans(fit2, ~ loc | bob_no2)

pairs(emm2)

\# To plot, first make data frame

emmdf <- as.data.frame(emm2)

\# And then actually plot

ggplot(emmdf, aes(x = bob_no2, y = emmean, color = loc, group = loc)) +

geom_line(size = 1.2) +

geom_point(size = 2.5) +

geom_errorbar(aes(ymin = lower.CL, ymax = upper.CL), width = 0.1) +

scale_color_viridis(discrete = TRUE, option = "C") +

labs(

x = "Head bob number",

y = "Predicted log head bob duration",

color = "Location"

) +

theme_classic() +

theme(

axis.title.x = element_text(size = 16),

axis.title.y = element_text(size = 16),

axis.text.x = element_text(size = 14),

axis.text.y = element_text(size = 14)

)

###################################################################

\#### Fig. 3: Total number of head bobs ############################

###################################################################

\# Data is skewed, log transform

hist(totbob$nr_bobs)

totbob$nr_bobslog = log(totbob$nr_bobs)

hist(totbob$nr_bobslog)

\# What Bels et al. 2025 did

bfit3 <-lmer(nr_bobslog ~ 0 + spp + (1|ID), data= totbob)

anova(bfit3) # result from Bels: F2,13 = 346, P < 0.0001

ranova(bfit3) # result from Bels: LRT, X2 = 5.8, df = 1, P = 0.016

summary(bfit3)

\# What we did

fit3 <-lmer(nr_bobslog ~ 0 + loc + (1|ID), data= totbob)

fit31 <-lmer(nr_bobslog ~ 0 + species + (1|ID), data= totbob)

AIC(fit3, fit31, bfit3)

anova(fit31)

ranova(fit31)

summary(fit31)

\# Post hoc model evaluation and predictions. Adapt to desired model

pr <- ggpredict(fit31, c("species"))

plot(pr)

z<-augment(fit31)

head(z)

hist(z$.resid)

shapiro.test(z$.resid)

plot(DHARMa::simulateResiduals(fit31))

\# Pairwise comparison test

emm <- emmeans(fit31, ~ species)

pairs(emm)

#####################################################################

\#### Fig. 4: Total duration of head bobs ############################

#####################################################################

\# Data is skewed, so log transform

hist(totdur$tot_dur)

totdur$tot_durlog = log(totdur$tot_dur)

hist(totdur$tot_durlog)

\# What Bels et al. 2025 did

bfit4 <-lmer(tot_durlog ~ 0 + spp + (1|ID), data= totdur)

anova(bfit4) # Bels results: F2,32= 730, P < 0.0001

ranova(bfit4) # Bels results: LRT, X2 = 32.8, df = 1, P < 0.0001

summary(bfit4)

\# What we do

fit4 <- lmer(tot_durlog ~ 0 + loc + (1| ID), data = totdur)

fit41 <- lmer(tot_durlog ~ 0 + species + (1| ID), data = totdur)

gfit4 <- glmmTMB(tot_dur)

AIC(fit4, fit41, bfit4)

anova(fit41)

ranova(fit41)

summary(fit41)

\# Post hoc model evaluation and predictions. Adapt to desired model

pr <- ggpredict(fit41, c("species"))

plot(pr)

z<-augment(fit41)

head(z)

shapiro.test(z$.resid)

plot(DHARMa::simulateResiduals(fit41))

\# Pairwise comparison test

emm <- emmeans(fit41, ~ species)

pairs(emm)
