## Supplementary material for "Reanalysis of Bels et al. (2025) given complex anthropogenic hybridization in *Iguana delicatissima* and intraspecific diversity in *Iguana iguana*": File S2

**Supplementary file 2 of van den Burg and Thibaudier concerning identified discrepancies in Bels et al. (2025).**

We have observed a range of discrepancies among figures, supplementary datafiles and corresponding figures of Bels et al. (2025), the cause of which we do not know;

- Between the article’s Highlights, Figure 2 and 3: according to Figure 2, no non-native *Iguana* had a display action pattern (DAP) of nine head bobs, which is in contrast to Figure 3 that indicates at least a single DAP sequence consisted of nine head bobs. The highlights mention “DAP of non-native iguanas involves two long and 5–9 short head bobs.”, suggesting there should at least be a single DAP with 11 head bobs. The supplementary materials do not include a DAP sequence of more than eight head bobs.
- Between Figure 2 and 3: according to Figure 2, no *I. delicatissima* DAP had seven head bobs, in contradiction to Figure 3. Indeed, two DAP sequences of *I. delicatissima* individuals had seven head bobs according to the supplementary materials.
- Between Figure 3 and supplementary materials: according to Figure 3 at least a single *I. delicatissima* DAP was one head bob in total length, although the minimum DAP sequence length in the supplementary materials is two.
- Between supplementary files 1-s2.0-S0944200625000030-mmc2 (hereafter ‘file 2’, containing DAP duration) and 1-s2.0-S0944200625000030-mmc4 (hereafter ‘file 4’, containing individual headbob duration): file 4 contains data from 106 DAPs, in contrast to the 102 DAPs in file 2. From file 2, one DAP is missing from CA, and three are missing from MA.
- Between supplementary files 2 and 4: In file 4, the shortest time for a complete DAP sequence, calculated through the sum of individual headbobs per sequence, is 2.58 seconds (Ind. 39, Colibri). However, there are four DAPs in file 2 that are shorter than this (Ind. 44, MA). This is not possible, since the sum of head bobs from file 4 does not contain the interval between individual headbobs. As the ID numbers between the two files do not correspond, it is not possible to filter out individual mistakes.
