## Supplementary material for "Reanalysis of Bels et al. (2025) given complex anthropogenic hybridization in *Iguana delicatissima* and intraspecific diversity in *Iguana iguana*": Table S1

**Supplementary file 3 of van den Burg and Thibaudier showing results of comparisons of individual head bobs between the four study sites.**

**Supplementary table. Pairwise comparison of the duration of individual head bobs in a display action pattern between iguanas from the four different locations: CO (Colibri: *Iguana delicatissima*), CA (Carangaise: hybrid), MA (Malendure: hybrid), and GO (Gosier: non-native). Significant results are indicated in bold.**

| **Head bob Number** | **Contrast** | **Estimate** | **Df** | **P value** |
| --- | --- | --- | --- | --- |
| 1 | **CA - CO** | **-1.4082** | **88.4** | **<0.0001** |
|  | CA - GO | 0.0311 | 95.8 | 0.9951 |
|  | CA - MA | 0.1527 | 69.6 | 0.6654 |
|  | **CO - GO** | **1.4393** | **101.9** | **<0.0001** |
|  | **CO - MA** | **1.5609** | **66.2** | **<0.0001** |
|  | GO - MA | 0.1215 | 77.2 | 0.8236 |
| 2 | **CA - CO** | **-0.8754** | **88.4** | **<0.0001** |
|  | CA - GO | 0.0208 | 95.8 | 0.9985 |
|  | CA - MA | 0.2538 | 67.1 | 0.2295 |
|  | **CO - GO** | **0.8962** | **101.9** | **<0.0001** |
|  | **CO - MA** | **1.1292** | **63.0** | **<0.0001** |
|  | GO - MA | 0.2330 | 74.7 | 0.3451 |
| 3 | **CA - CO** | **-0.4709** | **92.6** | **<0.0001** |
|  | CA - GO | 0.2326 | 95.8 | 0.2842 |
|  | CA - MA | 0.3089 | 67.1 | 0.0999 |
|  | **CO - GO** | **0.7035** | **105.8** | **<0.0001** |
|  | **CO - MA** | **0.7798** | **65.5** | **<0.0001** |
|  | GO - MA | 0.0763 | 74.7 | 0.9468 |
| 4 | CA - **CO** | -0.1696 | 108.8 | 0.4117 |
|  | CA - GO | 0.3339 | 99.8 | 0.0595 |
|  | CA - MA | -0.3233 | 67.1 | 0.0782 |
|  | **CO - GO** | **0.5035** | **126.8** | **0.0003** |
|  | CO - MA | -0.1537 | 75.7 | 0.5806 |
|  | **GO - MA** | **-0.6572** | **77.6** | **<0.0001** |
| 5 | CA - CO | 0.0469 | 166.1 | 0.9828 |
|  | **CA - GO** | **0.4781** | **115.3** | **0.0042** |
|  | CA - MA | -0.1042 | 79.4 | 0.8775 |
|  | **CO - GO** | **0.4311** | **174.3** | **0.0077** |
|  | CO - MA | -0.1511 | 109.6 | 0.6730 |
|  | **GO - MA** | **-0.5822** | **85.9** | **0.0007** |
| 6 | CA - CO | -0.1152 | 264.0 | 0.8754 |
|  | CA - GO | -0.3191 | 158.6 | 0.1659 |
|  | CA - MA | -0.1970 | 209.0 | 0.7188 |
|  | CO - GO | -0.2038 | 229.0 | 0.5139 |
|  | CO - MA | -0.0817 | 271.6 | 0.9699 |
|  | GO - MA | 0.1221 | 188.8 | 0.9092 |
| 7 | CO - GO | -0.3221 | 452.1 | 0.1864 |
|  | All other combinations | NA | NA | NA |
| 8 | All | NA | NA | NA |
