## Supplementary material for "Reanalysis of Bels et al. (2025) given complex anthropogenic hybridization in *Iguana delicatissima* and intraspecific diversity in *Iguana iguana*": Table S2

**Supplementary file 4 of van den Burg and Thibaudier showing results of the online survey of head bobbing male *Iguana iguana* from Clade II and IV (sensu van den Burg et al., 2026; Zootaxa, 5748, 387–399).**

| **Locality** | **Clade (sensu van den Burg et al., 2026)** | **Link** | **Long head bobs at beginning/end of DAP** |
| --- | --- | --- | --- |
| Guadeloupe (non-native population) | IV | Bels et al. (2025) | End |
| Suriname | IV | https://www.youtube.com/shorts/qps7sdecihQ | End |
| Suriname | IV | https://www.youtube.com/watch?v=gp02YhwYxwU | End |
| Suriname | IV | https://www.facebook.com/watch/?v=1276920360980203 | End |
| Venezuela | IV | https://www.shutterstock.com/es/video/clip-12837239-green-iguana?trackingId=1af14c88-bcbb-47b3-aa74-c60eed519702&listId=searchResults | End |
| Amazonia, Brazil | IV | https://www.tiktok.com/@ticksman/video/7502085076667223302?q=iguana%20amazonas&t=1770836831678 | End |
| Rondonópolis, Brazil | IV | https://www.tiktok.com/@marcielmanarotina/video/7508553216661998904?q=iguana%20brasil&t=1770837323681 | End |
| Manaus, Brazil | IV | https://www.tiktok.com/@userplanetaverdemauricio/video/7481286015777262853?q=iguana%20manaus&t=1770844340982 | End |
| Nordeste, Brazil | IV | https://www.tiktok.com/@amantedanatureza14/video/7558918620525448459?q=iguana%20nordeste&t=1770844702741 | End |
| Maranhao, Brazil | IV | https://www.tiktok.com/@messiasmoreiralim/video/7395359310659800326?q=iguana%20maranhao&t=1770849473773 | End |
| Acacias, Colombia | IV | https://www.tiktok.com/@janethtejeroofici/video/7457356645316693253?q=iguana%20meta%20colombia&t=1770839680669 | End |
| Venezuela | IV | https://www.tiktok.com/@eddissoneventos/video/7366995227325795590?q=bolivia%20iguana&t=1770846018441 | End, maar ziet alleen de laatste bob |
| Isla Margarita, Venezuela | IV | https://www.tiktok.com/@gabrielaandreina/video/7574209837563792696?q=iguana%20ciudad%20boliviar&t=1770848067713 | End |
| Onduidelijk | IV | https://www.tiktok.com/@yeyita_/video/7025788894608198917 | Einde |
| Panama (Flamenco island) | II | Dugan (1982) | Begin |
| “Colombia” | II | Distel and Veazey (1982) | Begin |
| Colombia | II | https://www.shutterstock.com/es/video/clip-3887628555-green-iguana-climbs-tree-branch-this-reptile?trackingId=2d2119fe-3616-47f2-973b-539ca8506a1e&listId=searchResults | Begin |
| Colombia | II | https://www.shutterstock.com/es/video/clip-25542485-iguana-sun-colombia?trackingId=741b2415-3778-46da-8efd-22729c415afb&listId=searchResults | Begin |
| Giron, Colombia | II | https://www.tiktok.com/@iguanaman_giron/video/7558101303491169592?q=iguana%20colombia&t=1770836248648 | Begin |
| Medelin, Colombia | II | https://www.tiktok.com/@chechoreina1/video/7562565816185556232?q=iguana%20colombia&t=1770836248648 | Begin |
| Colombia | II | https://www.tiktok.com/@mich.2580/video/7545693547781950726 | Begin |
| Colombia | II | https://www.tiktok.com/@noticiascaracol/video/7335572662728592645?q=iguana%20meta%20colombia&t=1770839680669 | Begin |
| Santa Marta, Colombia | II | https://www.tiktok.com/@camilo.gutierrezc/video/7350016701230009605?q=iguana%20caracas&t=1770846550325 | Begin |
| Barrancabermeja, Colombia | II | https://www.tiktok.com/@arbeyleon1111/video/7564808488166247701?q=iguana%20caracas&t=1770846550325 | Begin |
| Costa Rica | II | https://www.tiktok.com/@mmaureensolissala/video/7239515613029321990 | Begin |
| Costa Rica | II | https://www.tiktok.com/@ticojulius/video/7342701473484082437?q=iguana%20costarica&t=1770838303090 | Begin |
| Costa Rica | II | https://www.tiktok.com/@funsizedgirl69/video/7601641429290519829?q=iguana%20costarica&t=1770838303090 | Begin |
